# Differentiation of Mesenchymal Stem/ Stromal Cells into CD45+ Macrophage-like Cells: Expanding Insights into MSC Plasticity

**DOI:** 10.1101/2024.12.06.627092

**Authors:** Robert M. Rusch, Yo Mabuchi, Satoru Morikawa, Yoko Ogawa, Shigeto Shimmura

## Abstract

Mesenchymal stem/stromal cells (MSCs) are notable for their differentiation potential and immunomodulatory properties. They share several features with macrophages that are also similar to those of fibrocytes. This study aimed to investigate the relationship between MSCs, fibrocytes, and macrophages, focusing on the hypothesis of a possible direct lineage. This study focuses on a freshly isolated murine MSC population known as PαSmMSCs (CD45^−^ TER119^−^ Sca-1^+^ PDGFRα^+^). Transplanting PαSmMSCs from transgenic GFP mice into irradiated recipients produced GFP^+^ cells expressing the macrophage markers CD45, CD68, and CD115, indicating a macrophage-like phenotype. Further analysis showed that PαSmMSCs cultured in macrophage differentiation media acquired characteristics of M2 macrophages (CD45, CD68, and CD206). These results were confirmed in two different mouse strains as well as in human MSCs (hMSCs). Single-cell RNA sequencing revealed macrophage-like and fibrocyte-like populations among the PαSmMSCs. These findings demonstrate the phenotypic plasticity of freshly isolated MSCs to transition into macrophage-like and/or fibrocyte-like states, which has not been reported, particularly in long-term cultured MSCs. This study provides new insights into MSC plasticity and its potential roles in immune regulation and tissue repair.

## INTRODUCTION

Mesenchymal stem/stromal cells (MSCs) are widely used in medical applications owing to their immunomodulatory properties, differentiation potential, and low immunogenicity.[1, 2] Murine MSCs (mMSCs) are identified as CD45^−^ TER119^−^ Sca-1^+^ PDGFRα^+^ (PαSmMSC) or CD45^−^ TER119^−^ CD31^−^ CD73^+^ (CD73^+^mMSC).[3–5] Both populations are believed to overlap as they can both differentiate into various cells, including adipocytes, chondrocytes, osteoblasts, and myofibroblasts.[6, 7]

MSC-derived fibroblasts are vital for tissue homeostasis,[8] In contrast, Gli1^+^ MSC-derived myofibroblasts contribute to bone marrow failure through fibrosis,[9] with bone marrow-derived MSCs implicated in excessive myofibroblast accumulation and extracellular matrix deposition in various fibrotic diseases.[10, 11] While both cell types are linked to collagen production, several studies suggest that MSCs can synthesize collagen without differentiation,[12, 13] a capability exhibited by other cells such as fibrocytes.[14]

MSCs and fibrocytes are mesenchymal cells that are involved in wound healing, tissue repair, and fibrotic diseases.[15–20] As hematopoietic derivatives, fibrocytes express markers such as CD11b, CD14, CD34, and, most importantly, CD45[21] and migrate to injury sites where they transition to a mesenchymal phenotype and contribute to tissue fibrosis.[22]

Fibrocytes and macrophages have several similarities despite a distinct physical phenotype. Both have been associated with fibrosis, are hematopoietic-derived, and share several surface markers, leading some to suggest a lineage connection.[23, 24] Fibrocytes are most commonly defined as CD45^+^, CXCR4^+^, and Collagen I^+^, with the latter being the key marker distinguishing them from macrophages.[25–28] However, studies have shown that macrophages take up and degrade collagen, resulting in collagen positivity on immunohistochemistry.[29, 30] In addition, common macrophage markers, such as F4/80 or CD68, are found in fibrocytes.[28] Pilling et al. reported that only CD115 is exclusive to macrophages.[28]

The interactions between immune cells and MSCs have been extensively studied. Their immunomodulatory potential makes them promising candidates for treating immune-mediated diseases.[31] Numerous studies have shown that these cells exert immunomodulatory functions mainly via interactions with immune cells such as T, B, and natural killer cells.[32–34] Their low immunogenicity and ease of availability make them excellent candidates for stem cell therapy.[1, 2, 35]

However, other studies have suggested that MSCs can aggravate the disease. A study demonstrated that fresh (not cultured) MSCs contribute to the development of graft-vs-host disease.[36] Similarly, human bone marrow-derived MSCs upregulate major histocompatibility complex (MHC) class II expression when exposed to lower levels of IFNγ, acting as immunostimulants.[37] However, this effect was not observed at higher IFNγ levels. Another study showed that human MSCs can be polarized via toll-like receptors (TLR) signaling. MSCs also display a proinflammatory phenotype in underactivated immune systems. This polarization is mediated by TLR4 signaling, whereas the anti-inflammatory polarization is mediated by TLR3 signaling.[38] Owing to their similarities to monocytes, Waterman *et al.* suggested the terms MSC1 and MSC2, paralleling M1/M2 macrophages.

These bipolar characteristics resemble those of macrophages, which become pro- or anti-inflammatory in response to their environment.[39, 40] Pro-inflammatory MSCs also recruit lymphocytes and activate T cell secretion of macrophage inflammatory protein-1, CCL5, CXCL9, and CXCL10.[41–43] Anti-inflammatory MSCs suppress immune responses primarily when exposed to IFN-γ, TNF-α, and IL-1β,[44–46] with IFN-γ specifically, inducing PD-L1 and PD-L2 inhibitors.[32, 47] This characteristic is shared with bone marrow-derived macrophages, as IFN-γ can induce PD-L1 expression in tumor-associated macrophages, which promotes M2 polarization.[48, 49]

M1 or M2 macrophages are generally obtained by stimulating bone marrow cells with granulocyte-macrophage colony-stimulating factor (GM-CSF) for M1 or macrophage colony-stimulating factor (M-CSF) for M2 macrophages.[50–52] Other cytokines, such as IL-10, are often supplemented to enhance differentiation.[50, 52] On the other hand, MSCs are more heterogeneous and can undergo drastic changes when they are cultured.[53–55] Thus, cultured MSCs differ significantly from freshly isolated MSCs. Nevertheless, most studies used cultured MSCs because of their stability and ease of expansion compared to obtaining fresh MSCs.

Given the similarities among MSCs, fibrocytes, and macrophages, we aimed to investigate the relationship between these cells. We hypothesized that they could be directly linked through differentiation. To this end, we placed particular emphasis on using fresh MSCs instead of cultured MSCs since the latter may lack the heterogeneity of MSCs *in vivo*.

## RESULTS

### Freshly isolated mouse MSCs to transition into hematopoietic lineage

To validate the quality of isolated PαSmMSCs, cells were plated on a glass slide, and a colony-forming unit-fibroblasts (CFU-Fs) test was performed. The cells exhibited a characteristic spindle shape and formed colonies within a week (Supplemental Figure 1A). In addition, cells were isolated and investigated for possible CD45 contamination. TaqMan quantitative polymerase chain reaction (qPCR) revealed no contamination of the isolated samples (Supplemental Figure 1B).

To investigate the relationship between mMSCs, macrophages, and fibrocytes, we transplanted GFP^+^ PαSMSCs to irradiated mice. After 3 weeks post-transplantation, the lungs, lacrimal glands, and spleen were examined. These organs are involved in immune responses and associated with various diseases, including autoimmune disorders such as graft vs. host disease and Sjögren’s syndrome, which may be suitable for MSC therapy.[56, 57]

In all examined tissues, the presence of GFP^+^ CD45^+^ CD115^+^ CXCR4^+/−^ cells was observed in C57BL/6 (**Figure 1A**, **Figure 2A**) and B10.D2 (**Figure 1B**, **Figure 2B**) mice following transplantation of GFP^+^ PαSMSCs. No discernible or consistent patterns of cell distribution were observed within the tissues. In addition, the number of GFP^+^ cells varied significantly among the samples; however, they were generally scarce, except in the spleen.

**Figure 1.**
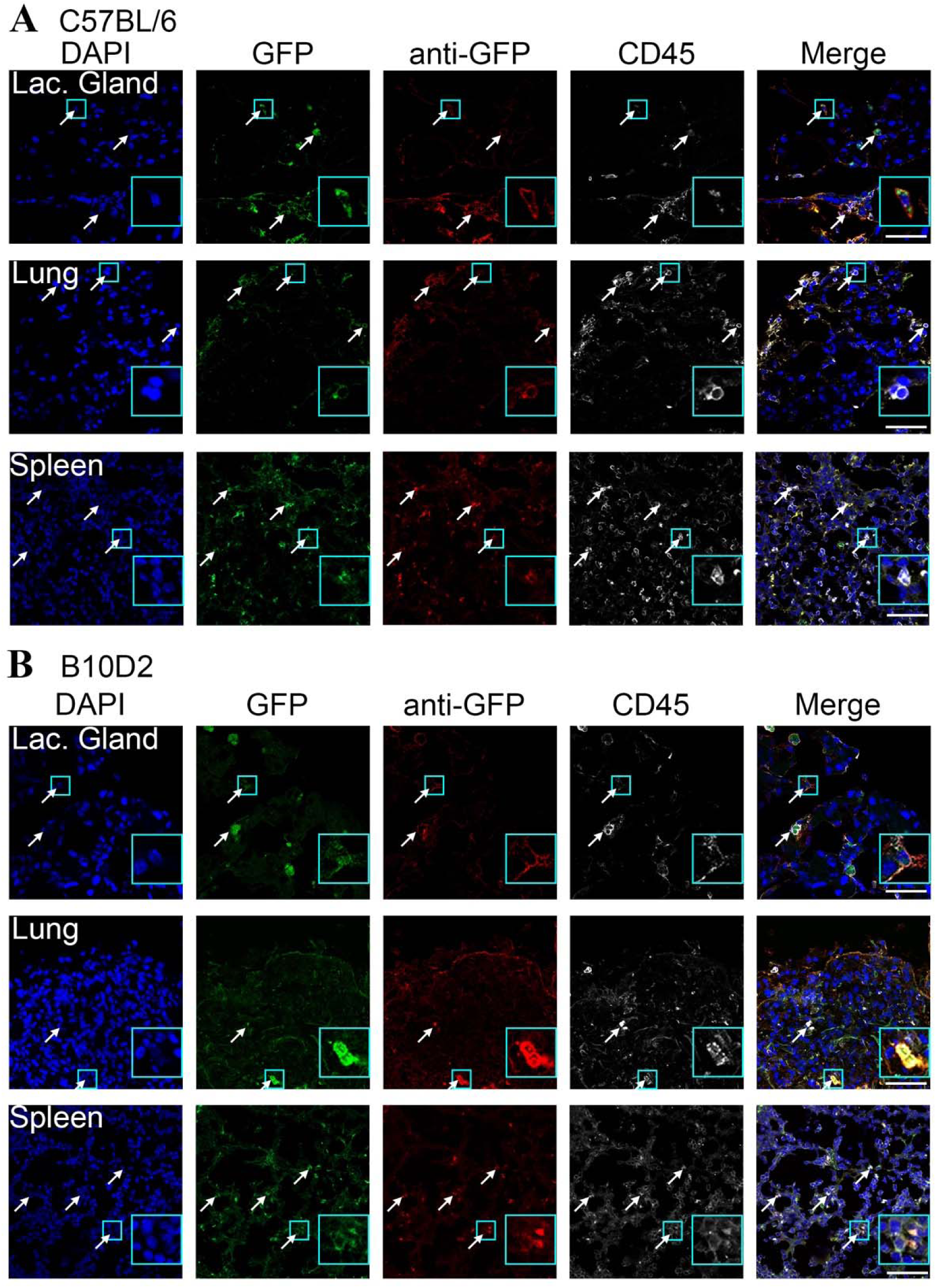
Murine MSCs (PαSmMSCs) from GFP mice were transplanted into irradiated C57BL/6 (A) or B10D2 (B) mice. A) GFP was observed in the lacrimal gland and lungs and verified using an anti-GFP antibody (red). CD45 (gray) was also expressed in GFP^+^ cells. B) Similar to A, GFP^+^ cells occasionally expressed CD45, indicating that this phenomenon was strain-independent. Scale bar = 200 µm.

**Figure 2.**
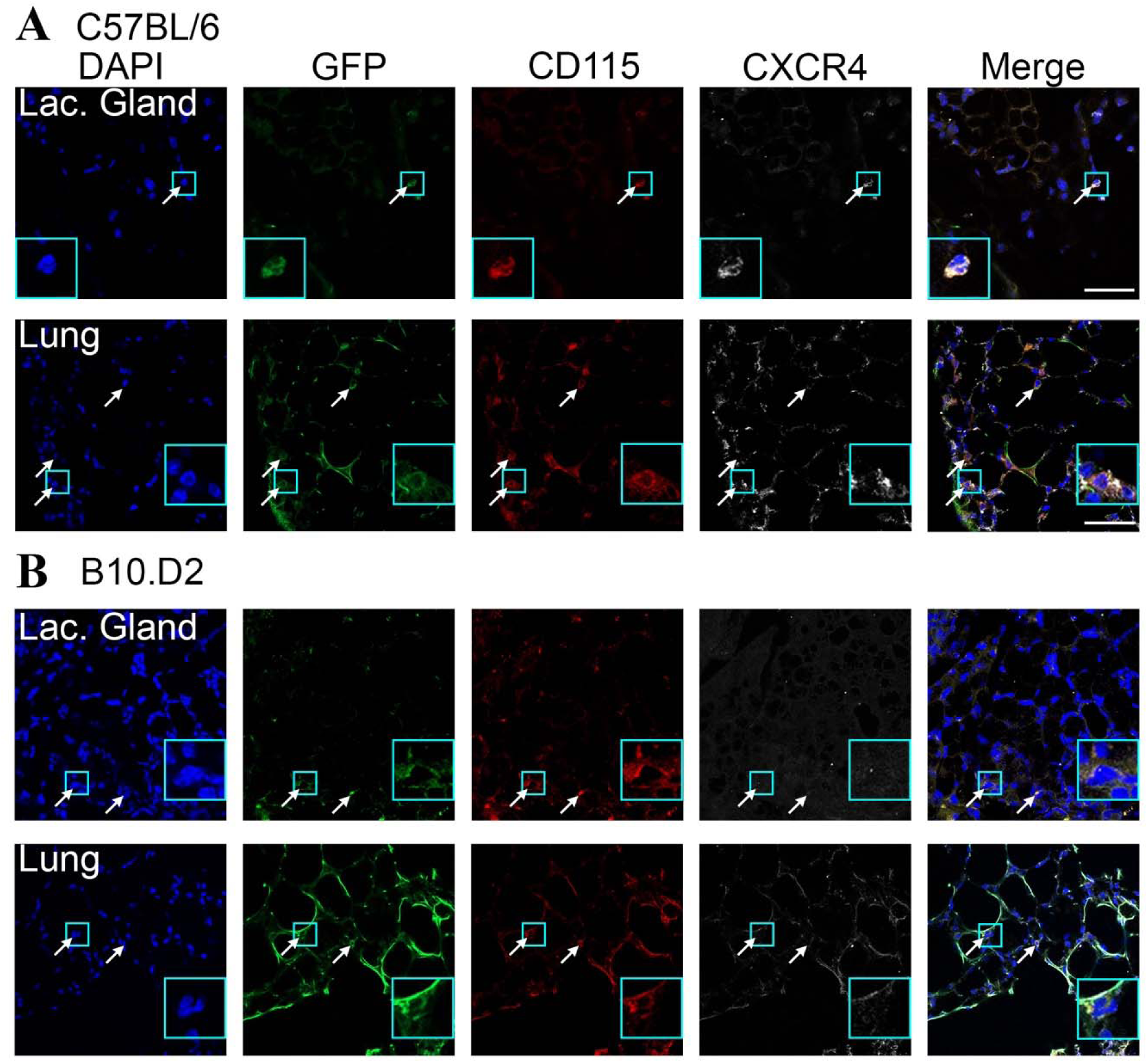
Transplanted GFP^+^ mMSC cells co-expressed CD115 (red) and CD45 (grey) in the lacrimal gland and lungs of both mouse strains. Scale bar = 200 µm.

To validate these observations, bone marrow and blood samples were collected and analyzed for the fibrocyte markers CD45 and CXCR4, as well as the macrophage marker CD115, which is not expressed by fibrocytes.[28] Flow cytometry revealed that many GFP^+^ cells in the blood (**Figure 3A, C**) and bone marrow (**Figure 3B, D**) were positive for CD45, CXCR4, and CD115 in both mouse strains. Notably, there were significantly more CD45^+^ GFP^+^ cells in the bone marrow than in the blood in both mouse strains (**Figure 3E**). Further subset analysis revealed that GFP^+^ CD45^+^ CXCR4^+^ CD115^+^ cells were dominant, consisting of 87% (C57BL/6) and 93% (B10.D2) of GFP^+^ CD45^+^cells in the blood (**Figure 3F**) and 96% (C57CL/6) and 86% (B10.D2) in the bone marrow (**Figure 3G**). The second most common subset was GFP^+^ CD45^+^ CXCR4^−^ CD115^+^ in both cases. In addition, blood-derived GFP^+^ cells formed a well-defined, distinct population of cells in the dot plot compared to the bone marrow.

**Figure 3.**
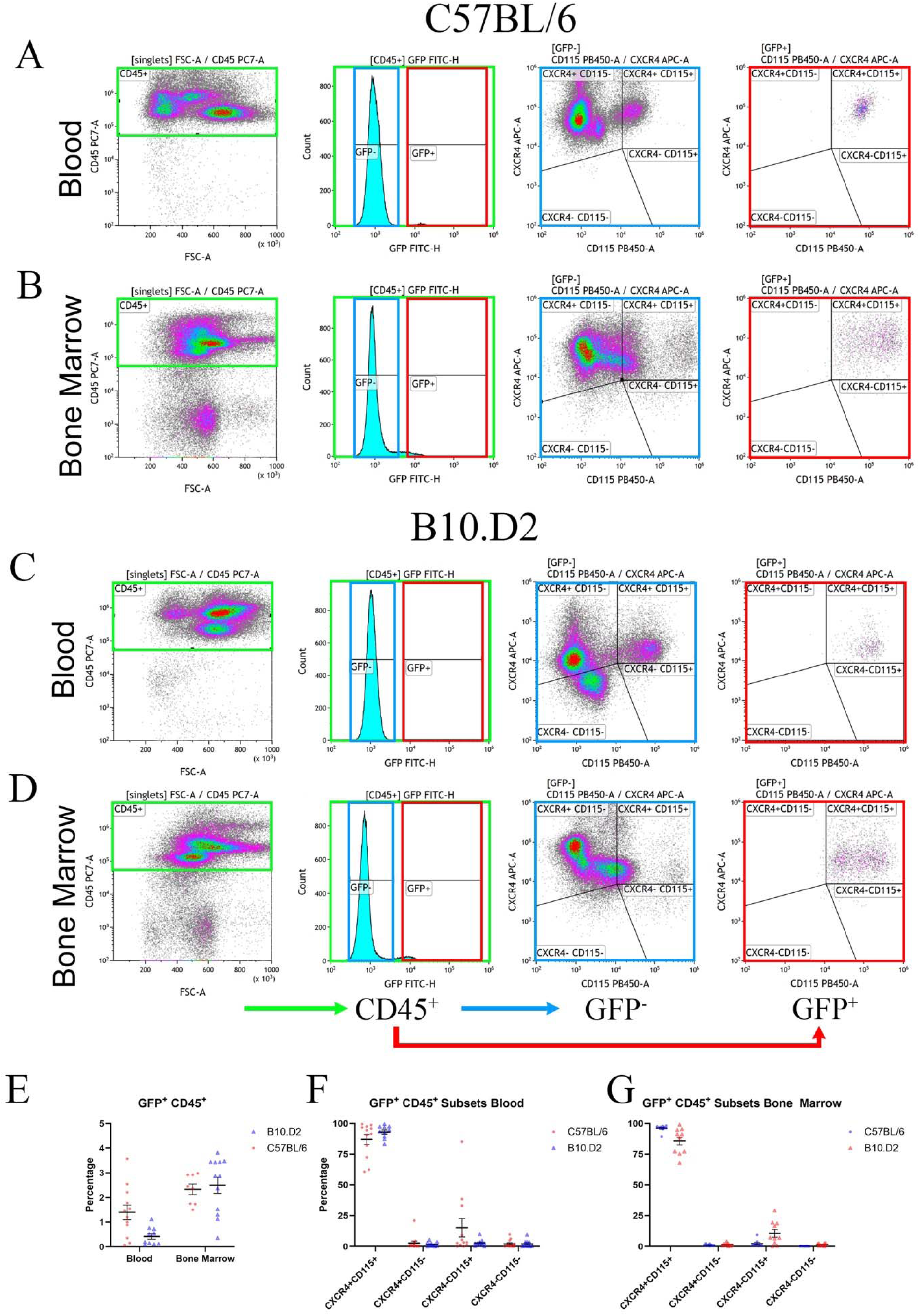
Flow cytometry analysis of blood samples. GFP^+^ mMSCs were isolated and transplanted to irradiated mice. Blood (A, C) and bone marrow (B, D) from C57BL/6 (A, B) and B10.D2 (C, D) mice were isolated and analyzed. Single cells were gated on CD45^+^ and GFP^+/−^ (E). The populations were characterized by CD115 and CXCR4 expression (F). A) In B10.D2 blood, most CD45^+^ and GFP^+^ cells co-expressed CXCR4 and CD115. B) Similarly, in the B10.D2 bone marrow, CD45^+^ and GFP^+^ cells predominantly expressed CXCR4 and CD115. However, the population appeared less condensed than that in the blood. C) In C57BL/6 blood, as in the B10.D2 samples, most GFP^+^ cells co-expressed CXCR4 and CD115. D) In the C57BL/6 bone marrow, the CD45^+^ GFP^+^ population was also CXCR4^+^ CD115^+^, although less densely clustered, similar to the pattern observed in B10.D2 bone marrow (B). E) CD45^+^ GFP^+^ cells were present in both samples and species, with more cells detected in the bone marrow than in the blood in both species. Blood exhibited a 1.4% (C57BL/6) and 0.4% (B10.D2) less presence of GFP^+^ cells compared to bone marrow, with 2.5% (C57BL/6) and 2.5% (B10.D2) within the CD45^+^ cells. F) The subset of CD45^+^ GFP^+^ CXCR4^+^ CD115^+^ cells was the most abundant, with 87% (C57BL/6) and 93% (B10.D2), and CD45^+^ GFP^+^ CXCR4^−^ CD115^+^ being the second most abundant, with 15% (C57BL/6) and 2.9% (B10.D2). The CD45^+^ GFP^+^ CXCR4^+^ CD115^+^ cell subset was also the most predominant in the bone marrow, comprising 96% of C57BL/6 and 86% of B10.D2 mice. In contrast, the CD45^+^ GFP^+^ CXCR4^−^ CD115^+^ subset represented the second largest group, accounting for 2.4% in C57BL/6 and 10.8% in B10.D2 mice.

### MSCs express macrophage markers upon stimulation with defined factors

To further investigate the expression of macrophage markers on mMSCs, fresh PαSMSCs were cultured in a macrophage differentiation medium. This classically involves M-CSF, also known as CSF1, with the corresponding receptor being CSF1R, also known as CD115 [58]. After isolation, the cells were plated on round cover glasses in a 12-well plate. The cells were cultured for 7 days in four different media: Dulbecco modified eagle’s medium (DMEM), Roswell Park Memorial Institute medium (RPMI), RPMI media supplemented with M-CSF (RM), M-CSF and IL-10 (RMI), and GM-CSF (RGM). These supplemented RPMI media are commonly used for bone marrow macrophage differentiation.[52, 59–61]

First, to assess CD45 expression, cells were stained with monoclonal and polyclonal antibodies against CD45 and the macrophage marker CD68. After 7 days, CD45 expression was not observed in DMEM (**Figure 4**). CD45 or CD68 expression was not observed in the RPMI medium as well. However, in RM and RMI media, a subset of PαSMSCs expressed CD45, of which most cells co-expressed CD68. Moreover, their cell morphology was clearly distinct from that of original PαSmMSCs, often with pseudopodia and sometimes with an elongated, spindle, or stellate morphology. Often, these cells formed uniform colonies composed solely of CD45^+^ cells; however, they also grew in mixed colonies alongside CD45^−^ cells.

**Figure 4.**
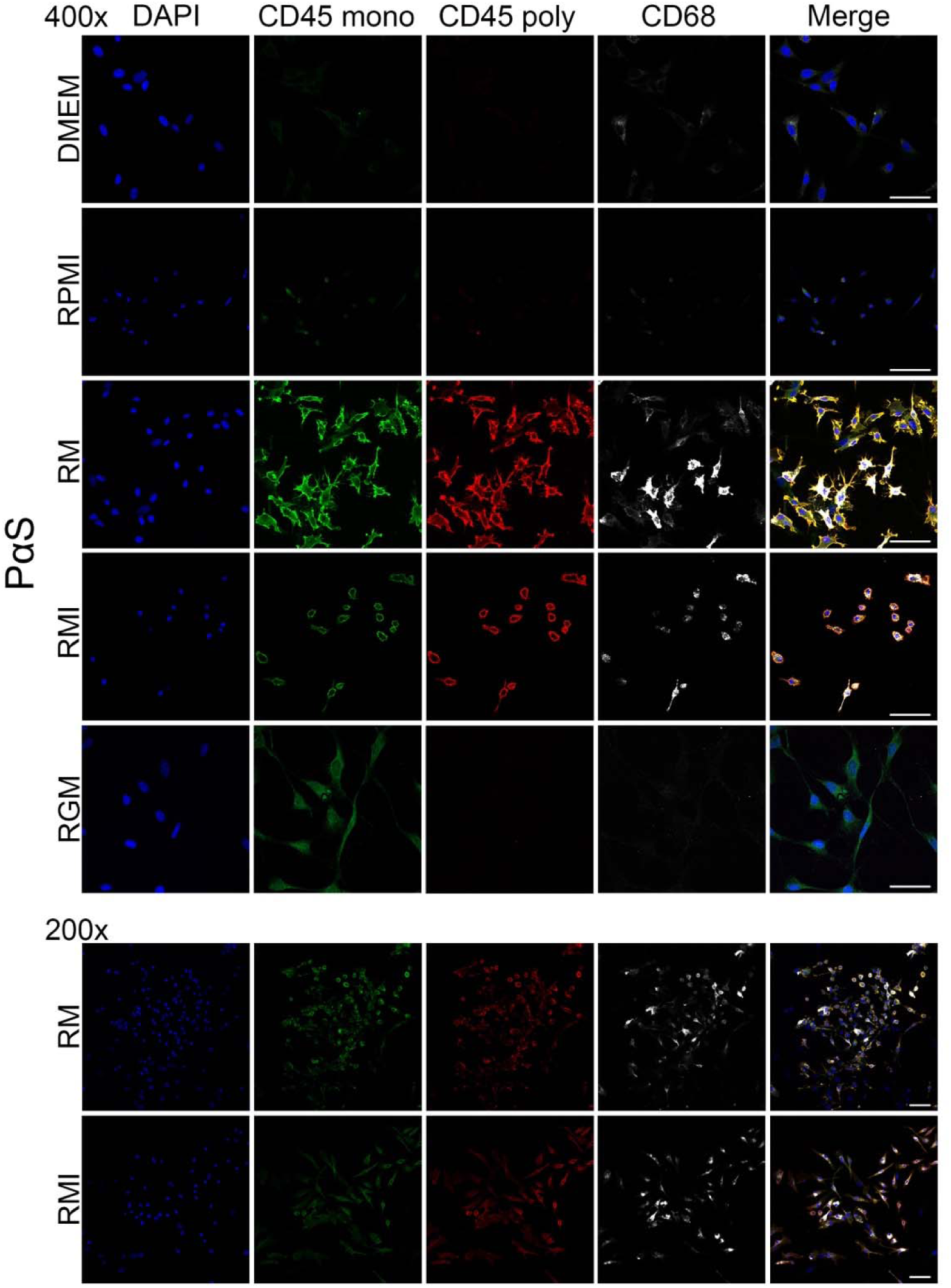
IHC staining of fresh PαSmMSCs cultured in various media for 1 week. A, B) cells in DMEM or RPMI displayed the typical spindle shape. C, D) When supplemented with M-CSF (C) or a combination of M-CSF and IL-10 (D), a subset of cells exhibited a shift towards a more irregular morphology, prominently characterized by pseudopodia. These cells expressed the markers CD45 (monoclonal antibody in green, polyclonal in red) and CD68 (gray). Although these morphological changes were observed in only a minority of the total cell population, areas with such cells showed a notable concentration. In these regions, the pseudopodia-bearing cells were occasionally intermixed with spindle-shaped CD45-negative cells or located adjacent to them. The addition of IL-10 did not have a significant impact. Scale bar = 200 µm.

Next, PαSmMSCs were stained for CD206 to distinguish between M1 and M2 macrophage phenotypes (**Figure 5**). Most (approximately 99%) cells that expressed CD45 and CD68 were also positive for CD206. These results were replicated with CD73^+^mMSCs, where they expressed CD45 and CD68 upon exposure to RM or RMI (**Figure 6**). The morphology was different compared to CD45^−^ cells and CD45^+^ CD68^+^ cells were also positive for CD206, as shown with PαSmMSCs. As a control, hematopoietic stem/progenitor cells (HSPCs) were isolated and cultured using the same protocol. However, after 1 week, no cells were observed (Supplemental Figure 3).

**Figure 5.**
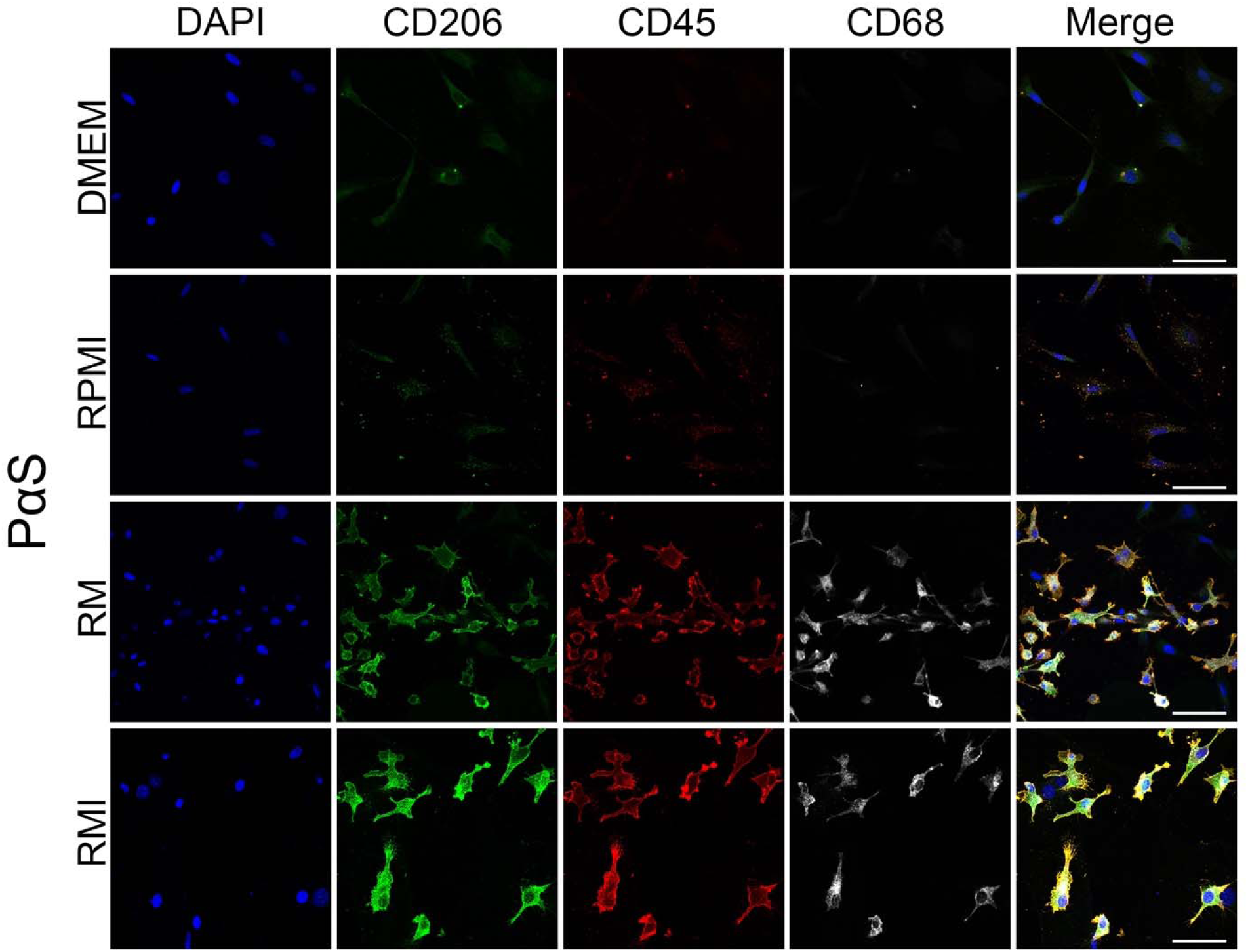
mMSCs were isolated and cultured in various media for 7 days. When adding M-CSF, cells expressed CD45 (red), CD68 (gray), and CD206 (green). No significant difference was observed when adding IL-10. Cells cultured in media lacking M-CSF showed a typical spindle-shaped appearance. Scale bar = 200 µm.

**Figure 6.**
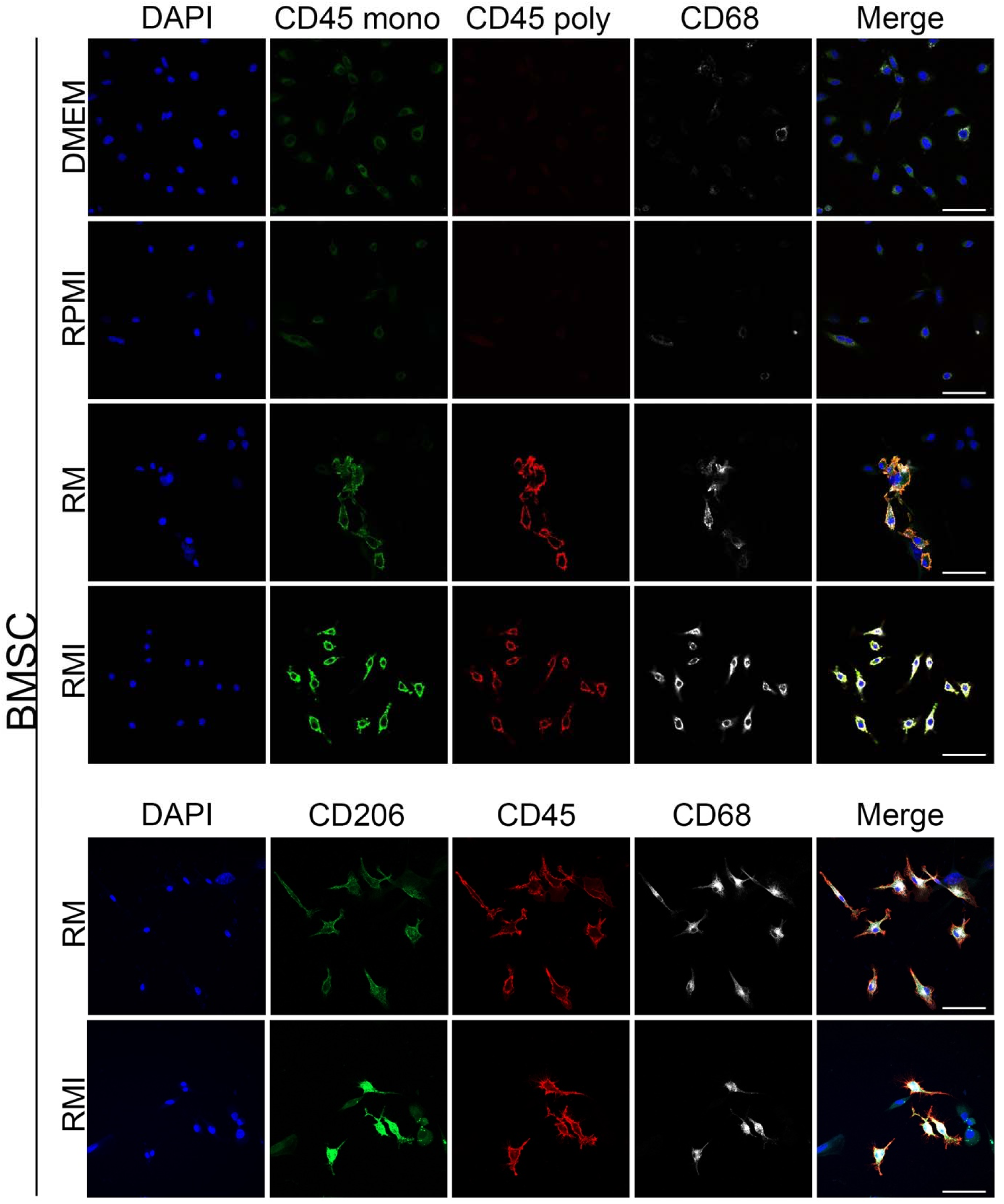
CD73^+^ mMSCs were isolated and seeded on glass slides in various media for 7 days, similar to PαSmMSCs. Cells were stained for CD206 (green), CD45 (red), and CD68 (gray). While no changes were observed in the DMEM or RPMI medium, media containing M-CSF had a significant yet varying number of CD206^+^ CD45^+^ CD68^+^ triple-positive cells. IL-10 supplement did not cause an apparent difference.

### Clustering of transitioned hematopoietic lineage cells by single-cell RNA-sequence

To further investigate these results, single-cell RNA-Seq (scRNA-seq) was performed with freshly isolated PαSmMSCs, which were cultured for 1 week in DMEM or RM. First, the t-distributed stochastic neighbor embedding was investigated for the positive isolation markers PGFRα and Sca-1 (ATXN1) (**Figure 7A**). Although both samples displayed a similar distribution, only the RM-cultured sample clearly contained CD45 (PTPRC)-expressing cells.

**Figure 7.**
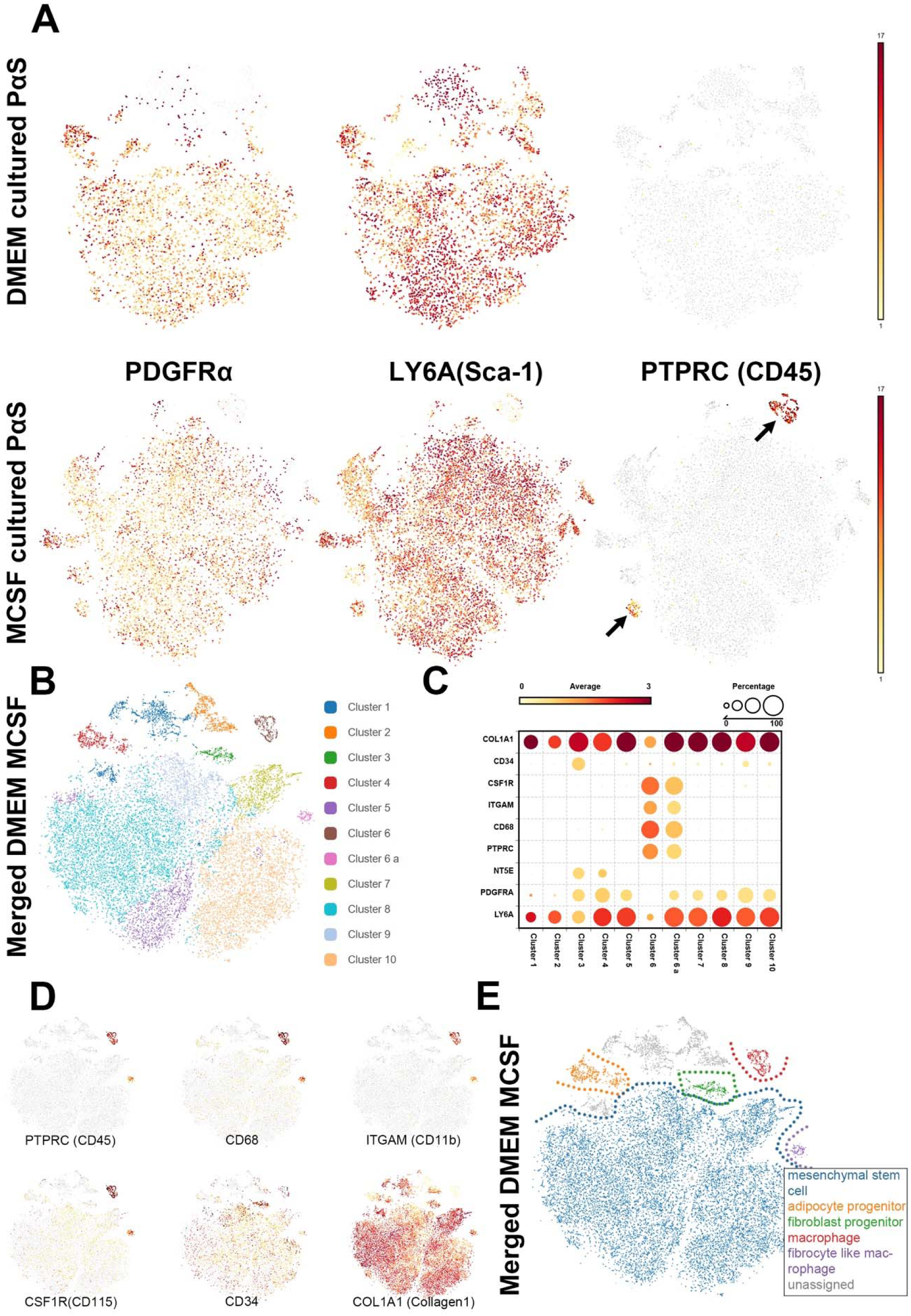
PαSmMSCs isolated as PDGFRα^+^, Sca-1^+^, CD45^−^, and TER119^−^ cells. A) When comparing PαSmMSC cultured in DMEM to PαSmMSCs cultured in M-CSF-containing RPMI medium on a single cellular level, the M-CSF-containing medium clearly showed two clusters of CD45^+^ cells. B) The data were merged to directly compare the two datasets and automatically clustered into ten clusters. However, owing to the differences in CD45 expression and the distance between the clusters shown in A, cluster 6 was manually divided into clusters 6 (high CD45) and 6a (low CD45). C) Cluster comparison focusing on PαSmMSC and CD73^+^mMSC isolation markers LY6A (Sca-1), PDGFRα, NT5E (CD73), and PTPRC (CD45). In addition to CD45, other common macrophage markers, such as CD68, ITGAM (CD11b), CSF1R (CD115), CD34, and the fibrocyte marker COL1A1 were investigated. Although Clusters 6 and 6a were similar in expression, their expression levels were different. Cluster 6a also shared similarities with all the other clusters; however, cluster 6 lost most of its isolation markers and strongly expressed macrophage markers. D) Distribution of key markers that distinguish clusters 6 and 6a from others, as well as CD34 and collagen1, which both spread across the blots. E) Owing to the expression of key markers shown in C, clusters were grouped and categorized as follows: mesenchymal cells (blue), adipocyte progenitors (yellow), macrophages (red), and fibrocyte-like macrophages (purple).

Next, the cells were clustered into 10 clusters using the Louvain Clustering of BBrowser (**Figure 7B**). However, owing to the distinct location and different levels of key marker expression, cluster 6 was manually subdivided into clusters 6 and cluster 6a (**Figure 7B, C**). The isolation markers Sca-1 (LY6A), PDGFRα (PDGFRA), CD73 (NT5E), CD45 (PTPRC), as well as CD68, CD11b (ITGAM), CD115 (CSF1R), CD34, and collagen 1 (COL1A1) were blotted in a bubble heatmap (**Figure 7C**). Clusters 4, 5, and 6a–10 all highly expressed Sca-1 and PDGFRα, whereas clusters 1, 2, and 3 expressed low levels of Sca-1 and PDGFRα, while cluster 6 did not express PDGFRα and expressed low levels of Sca-1. CD73 was only notably expressed in clusters 3 and 4. Cluster 3 was the only cluster with high CD34 expression.

While devoid in the other clusters, clusters 6 and 6a both expressed high levels of CD115, CD11b, CD68, and CD45. Notably, collagen 1 expression was high in all clusters except for cluster 6, which was approximately half of that in the others (**Figure 7C**).

Based on these markers and co-expression within clusters, cells were divided into MSCs (**Figure 7E**, blue, clusters 5, 7–10) based on Sca-1 and PDGFRα expression, adipocyte progenitors (yellow, cluster 4) based on CD73 expression, fibroblast progenitors (green, cluster 3) based on CD73 and CD34 expression, macrophages (red, cluster 6) based on CD45, CD68, CD11b, and CD115 expression, and fibrocyte-like macrophages (violet, cluster 6a) based on the low expression of macrophage markers and high expression of collagen I. Both macrophages and fibrocyte-like macrophages clearly express lineage markers such as CD14, CD48, CD53, STAB1, CCL9, and CD206 (MRC1), and lose characteristics of MSCs (**Figure 8**).

**Figure 8.**
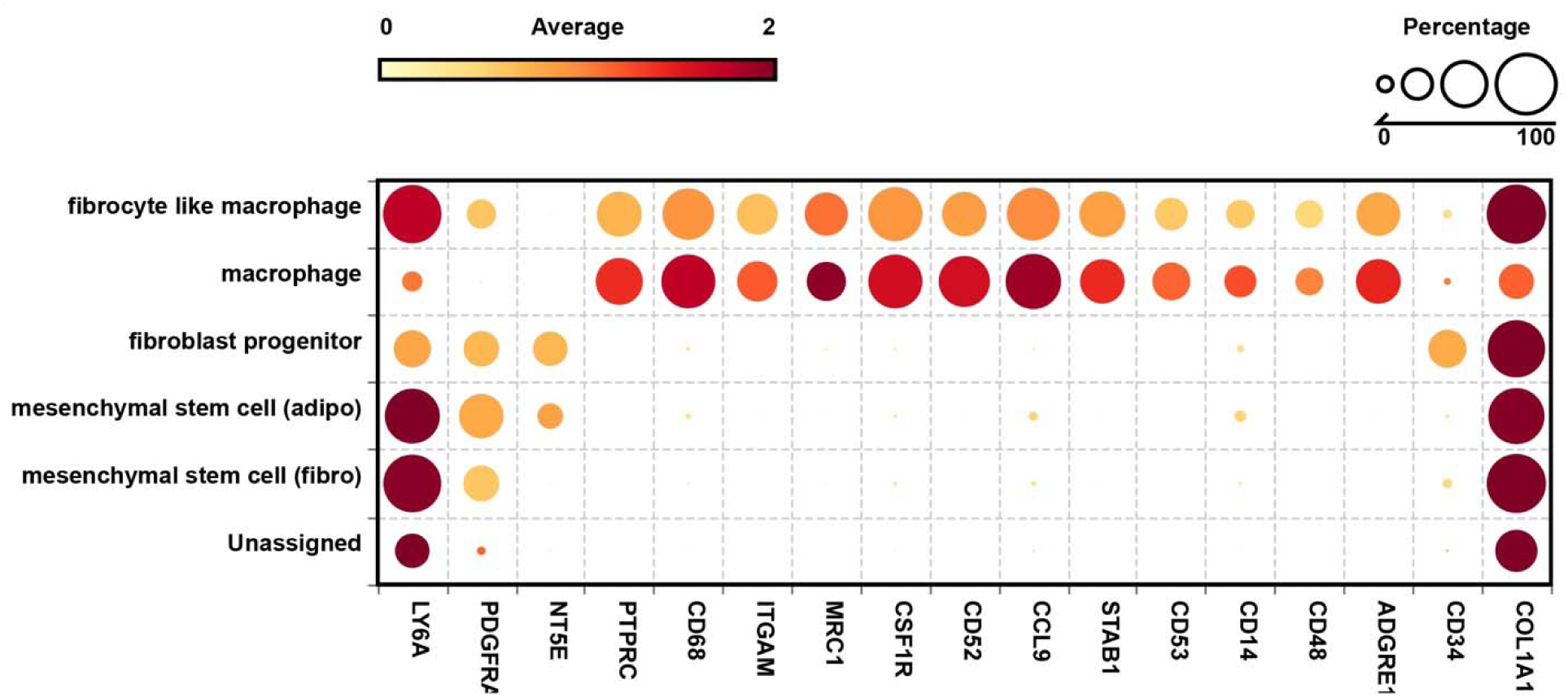
Bubble heatmap of the assigned cells for investigating the expression markers of the HSC lineage and specific macrophage markers. Apart from the previously mentioned markers, macrophages and fibrocyte-like macrophages also expressed CD206 (MRC1), CD52, CCL9, STAB1, CD53, CD14, CD48, and F4/80 (ADGRE1), further confirming their categorization.

### Freshly isolated human bone marrow MSCs show a similar phenomenon

Cryopreserved human whole bone marrow was thawed, and MSCs were isolated as CD271^+^ CD90^+^ cells (hMSCs) and cultured under the same conditions as mMSCs. After 1 week, hMSCs demonstrated similar morphology and marker expression as PαSmMSCs, where a subset of cells expressed CD45, CD68, and CD206 (**Figure 9**).

**Figure 9.**
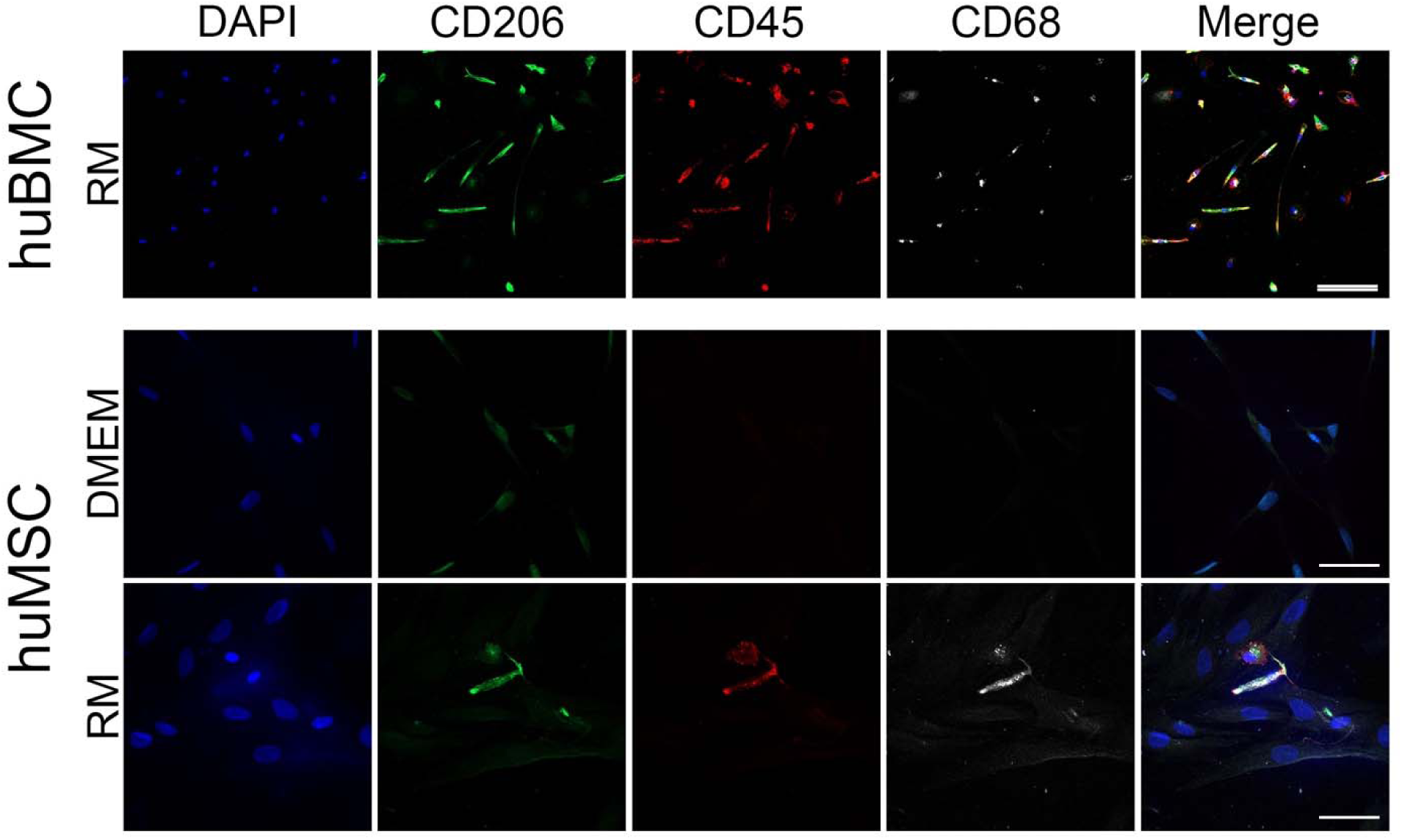
Immunohistochemistry staining was performed on human MSCs and bone marrow cells after 1 week of culture in M-CSF-containing medium. MSCs showed limited expression of CD206, CD45, and CD68 following exposure to M-CSF; however, their expression was relatively sparse.

## DISCUSSION

In this study, we determined the relationships between MSCs, fibrocytes, and macrophages. First, GFP^+^ PαSmMSCs from transgenic C57BL/6 mice were isolated and transplanted to irradiated C57BL/6 recipient mice. After transplantation, GFP^+^ cells became not only positive for CXCR4 and CD115, but also expressed CD45, a marker that is exclusively part of the hematopoietic lineage.[62] This result was reproducible with CD73^+^mMSCs using both, B10.D2 and C57BL/6 mice. Notably, GFP^+^ CD45^+^ CXCR4^+^ CD115^+^ cells were found in the blood and bone marrow, with the second most common type being GFP^+^ CD45^+^ CXCR4^−^ CD115^+^, both expressing the macrophage marker CD115. However, a strong preference for bone marrow was observed, suggesting that the bone marrow harbored the transplanted cells. The more distinct population detected in the blood by FACS suggests that cells enter circulation after maturation in the bone marrow. These results suggest that fresh mMSCs home to the bone marrow and differentiate into macrophage-like cells *in vivo*.

To further corroborate this *in vitro*, mMSCs and mHSPCs were cultured in two different macrophage differentiation media consisting of either M-CSF or GM-CSF, with the optional supplement IL-10. While DMEM or RPMI medium did not show changes in mMSCs, RPMI with M-CSF, as well as RPMI with M-CSF and IL-10 supplementation, changed the phenotype of the cultured mMSCs. In addition, these cells were CD45^+^ CD68^+^ CD206^+^, with the latter being a common marker of M2 macrophages. In the literature, M-CSF is often referred to as a key component of M2 macrophage differentiation, whereas GM-CSF is attributed to M1 differentiation.[63] However, when cultured in GM-CSF, no transition was observed in PαSmMSCs. No HSCs were present in any media. In contrast to MSCs, HSPCs do not adhere to glass and were most likely washed out.[64, 65] While host macrophages might become GFP^+^ by phagocytosing transplanted GFP^+^ PαSmMSC cells, the *in vitro* experiment ruled out this possibility.

In M-CSF-supplemented culture conditions, the number of CD45^+^ cells varied from a few single cells to large colonies where CD45^+^ cells coexisted next to CD45^−^ cells. Quantification is difficult when combined with varying amounts of plated MSCs. However, scRNA-seq data suggested that the percentage of CD45^+^ cells was approximately 5.85%. scRNA-seq was performed to confirm these results. A small population of CD45^+^ and CD68^+^ cells was identified, which was also positive for CD11b, CD206, and CD48, commonly expressed on various hematopoietic cell surfaces. Therefore, these cells were identified as macrophages. Notably, another small population exhibited a profile similar to that of macrophages; however, the expression levels of the aforementioned key markers were generally lower. Instead, the isolation marker PDGFRα, especially Sca-1 and collagen 1, were strongly expressed, indicating that these cells might be in between the MSC and macrophage phenotype and resemble fibrocytes. MSCs naturally share several markers with fibrocytes, such as CXCR4 and SMA,[66, 67] and M2 macrophages can produce collagen (adopt a fibrocyte feature) within atherosclerotic plaques,[68] further highlighting the similarities between these cells.

The expression of monocyte markers, such as CD11b, CD48, and CD68, signifies a macrophage phenotype.[69, 70] CD206 is a key marker for M2 macrophages, known for anti-inflammatory functions through TGF-β and IL-10, similar to MSCs [41, 71]. M2 macrophages contribute to wound healing in a manner similar to that of MSCs and recruit fibroblasts, a cell type “phenotypically indistinguishable from MSCs.”[72, 73] In contrast, MSCs can induce an M2-like phenotype in macrophage co-cultures, supporting a close relationship between these cells.[74]

This study also revealed a notable decrease in PDGFRα expression consistent with increasing macrophage markers amongst the populations. While the exact role of PDGFRα in macrophage function and behavior remains underexplored, its expression may not be significant in macrophages.[75]

Other research groups reported similar results. For example, CD49^+^ MSCs with CFU-F capability exhibited a CD34^+^ CD45^med/low^ phenotype after isolation, which was substantially downregulated during culture.[66] Another recent study found “distinct CD45^−^ B-lymphoid and erythroid progenitor populations, whose activity was enhanced by BM stromal cells” within CD45^−^ CD31^−^ Ter119^−^ cells. Notably, these triple-negative cells replaced over 95% of CD45^+^ cells in lethally irradiated mice 4 months after transplantation.[76] MSCs are traditionally seen as invisible immunoregulators; however, they can occur in the immune system through MHC II upregulation, suggesting a more direct role.[77] These studies have shown that macrophages, fibrocytes, and MSCs are closely associated. However, increasing evidence suggests that this relationship is not merely collaborative. The data presented in this study showed that mMSCs can change phenotype from CD45^−^ to CD45^+^ *in vivo* in multiple strains as well as *in vitro*.

This study showed that CD271^+^ CD90^+^ hMSCs showed the same reaction to M-CSF as demonstrated by PαSmMSCs and CD73^+^mMSCs, indicating that this behavior is generally applicable, as demonstrated in multiple strains and species. However, the number of isolated cells from human bone marrow was much lower compared to the murine model, which may explain the scarcity and lack of large CD45^+^ cell populations. Further refinement of culture protocol may enhance the size of CD45^+^ cell populations.

While it can be argued that MSC populations may have been contaminated with hematopoietic cells that subsequently became CD45^+^ cells, contamination was unlikely since the data were consistently replicated (N=8), with no CD45 expression in the unsupplemented medium. Furthermore, HSCs generally do not attach to uncoated glass, a feature commonly used to isolate MSCs from the bone marrow. In addition, qPCR showed no CD45 expression of the cells immediately after isolation.

To date, no study has investigated whether MSCs change their phenotype before performing their immunomodulatory roles. This study demonstrated that, in addition to CD45, macrophage markers CD11b and CD68, and to some degree fibrocyte markers such as CXCR4 and collagen I, were co-expressed in mMSCs both *in vivo* and *in vitro,* which was confirmed by single-cell RNA-seq. While it is established that MSCs can become fibroblasts, this study suggests that freshly isolated MSCs can develop a fibrocyte-like and macrophage-like phenotype. Currently, it is unclear whether there are two steps on the same path or two individual paths. This macrophage-like state is more specific to anti-inflammatory CD206^+^ M2 macrophages, which share many immunomodulatory features through the same mechanisms as MSCs, as demonstrated previously in numerous studies.

In conclusion, this study revealed that freshly isolated bone marrow MSCs are more representative of *in vivo* MSCs than conventionally cultured MSCs. We believe that our data convincingly show that MSCs have the potential to transition to what has hitherto been considered an HSC line. This highlights additional MSC potential and indicates the need to reevaluate the current understanding of MSCs.

## EXPERIMENTAL PROCEDURES

### Animals

The animals (C57BL/6NCrSlc, C57BL/6-Tg(CAG-EGFP), B10.D2/nSnSlc) were procured from Sankyo Labo Service Corporation (Tokyo, Japan). B10.D2/nSnSlc were bred with C57BL/6-Tg(CAG-EGFP) mice and inbred for over 10 generations to create B10.D2 EGFP mice. All mice were 8 weeks old at the time of bone marrow acquisition. Mice obtained through Sankyo Labo Service Corporation were female, bred transgeneic EGFP mice were mixed genders due to limits of availability. Animal studies were conducted in compliance with protocols approved by the Institutional Animal Care and Use Committee of Keio University (#A2022-178 and #A2022-122). These procedures strictly adhered to the ARVO Statement for the Use of Animals in Ophthalmic and Vision Research, and all experimental methods were performed in accordance with Keio University’s Institutional Guidelines on Animal Experimentation.

### MSC isolation

Bone marrow was obtained as previously described with minor modifications.[36]. Briefly, tibia, femur, and coxal bones were extracted and washed thrice with HBSS (Nacalai Tesque, Kyoto, Japan) containing 10 mM HEPES (Nacalai Tesque, Kyoto, Japan), fetal bovine serum (JRH Biosciences, USA), and 1% penicillin/streptomycin (Nacalai Tesque, Kyoto, Japan). Next, the bones were crushed and cut using scissors to obtain a paste-like consistency. The bones of 5–7 animals (roughly divided in half) were incubated with 20 mL 0.2% (g/v) collagenase (Wako, Osaka, Japan) containing DMEM/Ham’s F-12 (Nacalai Tesque, Kyoto, Japan) in a MACSmix™ Tube Rotator (Miltenyi Biotech, Bergisch Gladbach, Germany) at 37 °C for 1 h, with the addition of 10 µL DNase (Qiagen, Venlo, Netherlands) 15 min before the end (after 45 min).

After collecting the medium and washing the bones, the cells were centrifuged at 300 ×g for 5 min at room temperature, and blood cells were lysed using 1 mL water per tube (Nacalai, Kyoto, Japan). Bone marrow cells were then transferred to a 1.5 mL Eppendorf tube for antibody staining.

Per 1×10^7^ cells, 1 µL of [PE-Sca1 (clone: D7), PE-Cy7-CD45 (clone: 30-F11), PE-Cy7-TER119 (clone: TER-119), and 2 µL of APC-PDGFRα (clone: APA5)] or [PE-Cy7-CD31 (clone: MEC13.3), PE-Cy7-CD45 (clone: 30-F11), PE-Cy7-TER119 (clone: TER-119), and CD73 (clone: TY/11.8)] (all Biolegend, San-Diego, CA, USA) were added to cells in 100 µL HBSS wash buffer. After 30 min on ice in the dark, cells were washed and isolated using a BC MoFlo XDP (Beckman Coulter, Brea, CA, USA). MSCs were isolated as CD45^−^, TER119^−^, PGFR^+^, Sca-1^+^ (PαSmMSC) or CD31^−^, CD45^−^, TER119^−^, and CD73^+^ (CD73^+^mMSC).

### Isolation verification with TaqMan

To validate the isolation, cells were isolated as described above for PαSmMSC, and RNA was isolated using NucleoSpin® RNA XS (Takara, Shiga, Japan) according to the manufacturer’s protocol. Next, RNA was analyzed using TaqMan™ Fast Advanced Master Mix for qPCR according to the manufacturer’s protocol. The following primers were used: TaqMan™ Assay FAM-MGB PTPRC (CD45) (20X) (Assay ID: Mm01293577_m1) (Thermo Fisher, Waltham, MA, USA) and TaqMan™ Gene Expression Assay, VIC primer-limited (Assay ID: Mm99999915_g1).

CD45^+^ cells from the bone marrow, cultured MSC, and water (no RNA) were used as references and controls.

### CFU-F test

PαSmMSCs cells were plated onto a microscope cover glass in a 6-well plate and incubated under standard conditions to allow for colony formation. Next, the cells were washed with phosphate buffer saline (PBS) and fixed with 4% paraformaldehyde in PBS for 15 min at room temperature. After washing, a 0.1% boluidine blue (Sigma Aldrich, St. Louis, MO, USA) solution was added to the cells and incubated at room temperature for 15 min. After three washes, the cells were mounted on a microscope slide and analyzed using Bio.

### Bone marrow transplantation

Recipient mice (8-week-old) were irradiated at 7 Gy. Subsequently, 2 × 10^4^ GFP^+^ PαSmMSCs isolated from GFP transgeneic mice together with 1×10^5^ GFP^−^, CD45^+^, and TER119^+^ cells from wild-type mice were transplanted through the tail vein in 200 µL DMEM F12 (Nacalai, Kyoto, Japan).

### Immunohistochemistry

The tissues were fixed in a 10% buffered formalin solution for 3 h. Following fixation, samples were briefly rinsed and then immersed in OCT compound (Sakura Finetek Japan, Co., Ltd., Japan) before being frozen at −80 °C. Subsequently, frozen samples were sectioned using a cryostat.

Cell culture samples were each washed thrice with PBS (Nacalai, Kyoto, Japan) for 5 min and fixated with 4% formalin (Nacalai, Kyoto, Japan) for 15 min at room temperature. After three PBS washings, fixated cells were either stored in PBS at 4 °C or stained immediately.

Samples were blocked with Normal Goat Serum (Thermo Fisher, Waltham, MA, USA) or 10% Normal Donkey Serum (Sigma Aldrich, St. Louis, MO, USA) with 1% Triton X-100 (Nacalai, Kyoto, Japan) for 60 min at room temperature. GFP-containing samples were protected from light. Next, the samples were incubated overnight at 4 °C with the primary antibody (Table 1). The following day, the sections were washed three times with PBS and stained with Alexa Fluor 555- or 647-labeled secondary antibody for 30 min at room temperature. After washing three times, the cells were mounted with DAPI and anti-fading reagent-containing mounting medium (Abcam, Cambridge, UK). Images were obtained using a Leica LSM 980 microscope (Leica, Wetzlar, Germany).

**Table 1.** Antibody list for tissue sections and cell culture.

| Name | Clone | Reporter | Company | Dilution |
| --- | --- | --- | --- | --- |
| Anti-mouse CD45 | 30-F11 | PE | Biolegend | 1:500 |
| Anti-mouse CD45 | 30-F11 | APC | Biolegend | 1:500 |
| Goat anti-mouse CD45 | Polyclonal | None | R&D Systems | 1:500 |
| Goat anti-GFPuv | Polyclonal | None | R&D Systems | 1:500 |
| Anti-mouse CD68 | FA-11 | APC | Biolegend | 1:500 |
| Anti-mouse CXCR4 (flow cytometry) | 1276F12 | APC | Biolegend | 1:500 |
| Anti-mouse CXCR4 (IHC) | UMB2 | None | Abcam | 1:500 |
| Anti-mouse CD115 | AFS98 | None | Thermo Fisher | 1:500 |
| Donkey anti-Goat IgG | Polyclonal | Alexa Fluor® 555 | Invitrogen | 1:400 |
| Goat Anti-Rabbit IgG | Polyclonal | Alexa Fluor® 647 | Abcam | 1:400 |
| Anti-human CD45 | HI30 | PE | BioLegend | 1:500 |
| Anti-human CD206 | 15-2 | A488 | BioLegend | 1:500 |
| Anti-human CD68 | Y1/82A | APC | BioLegend | 1:500 |

### Flow cytometry

Blood and bone marrow samples were collected on the day of the acquisition. Before staining, the red blood cells were lysed using water (Nacalai Tesque, Kyoto, Japan). Bone marrow cells were cultured overnight on 6 cm FNC-coated (Athena Enzyme Systems, Baltimore, MD, USA) plastic dishes in DMEM/Ham’s F-12 (Nacalai, Kyoto, Japan). The following day, the supernatant was aspirated, and only the adherent cells were stained.

Initially, cells were washed with FACS buffer (R&D Systems, Minneapolis, MN, USA) thrice and centrifuged at 300 ×g and 4 °C for 5 min. Subsequently, the cells were stained with primary antibodies against APC-CXCR4 (1276F12), PE-Cy7-CD45 (30-F11), and BV421-CD115 (AFS99) (BioLegend, San Diego, CA, USA) for 30 min on ice. The cells were washed with the staining buffer and analyzed using BC CytoFLEX S (Beckman Coulter, Brea, CA, USA).

A gating strategy was developed using Fluorescence Minus One (FMO) controls and wild-type cells for GFP^−^ signaling. If the event count for GFP^+^ cells was <50 for blood or <100 for bone marrow, the data were excluded.

### Cell culture

Round cover glasses (18 mm; Matsunami, Osaka, Japan) were placed in a 12-well plate. MSCs from wild-type mice were isolated as previously described. Next, 1.0–1.5 × 10^4^ MSCs were seeded in individual wells, with each well receiving 1 mL of the medium listed in Table 2. The base media used were DMEM/Ham’s F-12 (Nacalai, Kyoto, Japan) or RPMI 1640 (Nacalai, Kyoto, Japan). After plating, the cells were cultured undisturbed for 2 days. From day 3, 500 µL were exchanged for fresh medium daily until day 7. On the last day, cells were fixed with 4% paraformaldehyde, as described above.

**Table 2.** Media composition for cell culture.

| Base medium | Abbreviation | FBS | P/S | Glutamax (Gibco) | M-CSF (Biolegend) | IL-10 (Biolegend) | GM-CSF (Biolegend) |
| --- | --- | --- | --- | --- | --- | --- | --- |
| DMEM | DMEM | 10% | 1% | - | - | - | - |
| RPMI | RPMI | 10% | 1% | 2 mM | - | - | - |
| RPMI | RM | 10% | 1% | 2 mM | 50 ng/mL | - | - |
| RPMI | RMI | 10% | 1% | 2 mM | 50 ng/mL | 25 ng/mL | - |
| RPMI | RGM | 10% | 1% | 2 mM | - | - | 50 ng/mL |

### RNAseq data acquisition

Cells were cultured in DMEM or RM for 1 week, as described above. Next, cells were washed and processed as described in the Chromium Single Cell 3’ Reagent Kits User Guide (v3.1 Chemistry Dual Index, 10x Genomics, Pleasanton, CA, USA). The sequencing platform used was the NovaSeq 6000 (Illumina, San Diego, CA, USA).

### HSPC isolation and culture

The femur and tibia were harvested and crushed using a pestle, as described above. Next, the whole bone marrow was stained with PECy7-Lineage [CD3ε (clone: 145-2C11), CD45R (clone: RA3-6B2), Gr-1 (clone: RB6-8C5), CD11b (clone: M1/70), Ter119 (clone: TER-119)], APC-cKit (clone: 2B8), FITC-Sca1 (clone: D7), and PE-CD45 (clone 30-F11) (all Biolegend, San-Diego, CA, USA). Cells were stained as described above and sorted into Lin^+^, c-Kit ^+^, and Sca1^+^ cells. After isolation, the cells were directly plated in a 12-well plate with 18 mm round cover glasses (Matsunami, Osaka, Japan) in DMEM, RPMI, RM, RMI, and RGM.

### RNA-seq data analysis

Data dimensionality reduction and visualization were conducted using BioTuring’s BBrowserX [78]. Principal Component Analysis, k-nearest neighbors, Venice binarizer, and t-distributed stochastic neighbor embedding (t-SNE) methods were used for the analysis. Visualization was achieved using the Vinci software. Louvain clustering was performed at a resolution of 0.5.

### Human MSC isolation

BONE MARROW MNC (POIETICS, Lonza, Basel, Switzerland) was suspended in ice-cold HBSS at 1– 5×10^7^ cells/mL and stained for 30 min on ice with a monoclonal antibody. The following antibodies were used: LNGFR (CD271)-PE (clone ME20.4-1. H4; Miltenyi Biotech, Bergisch Gladbach, Germany), and THY-1(CD90)-APC (clone 5E10; BioLegend, San Diego, CA, USA). Propidium iodide (PI: 2 μg/mL) was used to eliminate dead cells during flow cytometric analysis. Flow cytometric analysis and sorting were performed using a BD FACSAria (BD Biosciences, Franklin Lakes, NJ, USA).

### Statistical analysis

All statistical analyses were performed using GraphPad Prism 9. Statistical tests used throughout the study are indicated in the relevant figure legends and results sections. The primary statistical test used was the unpaired t-test. Displayed data represent means ± SEM. Significance was defined as follows: *p* < 0.05 (\**), p < 0.01 (**), and p < 0.001 (\*\*\**). Each experiment was conducted independently at least three times. Randomization and stratification were not applicable due to the nature of the experimental design. In flow cytometry data was excluded if the event count for the target population was below 50 (blood) or below 100 (bone marrow)

## ACKNOWLEDGMENTS

This work was supported by a grant from the Japan Society for the Promotion of Science (KAKENHI) 23K24504. We thank Hiroko Taniguchi for her technical assistance in the laboratory. We also acknowledge Mari Fujiwara at Keio University School of Medicine, Central Equipment Management Division, for her assistance with FACS cell isolation and for providing access to the FACS sorting machine.

## AUTHOR CONTRIBUTIONS

Conceptualization: R.M.R., Y.M., Y.O., S.S. Methodology: R.M.R., Y.M., Y.O. Validation: R.M.R. Formal Analysis: R.M.R. Investigation: R.M.R., Y.M., Y.O. Data Curation: R.M.R. Writing – Original Draft: R.M.R. Writing – Review & Editing: R.M.R., Y.M., S.S. Visualization: R.M.R. Resources: Y.M., S.M., S.S. Supervision: Y.O., S.S. Project Administration: R.M.R., Y.M., S.S. Funding Acquisition: S.S.

## DECLARATION OF GENERATIVE AI AND AI-ASSISTED TECHNOLOGIES

During the preparation of this work, the author(s) used ChatGPT in order to optimize grammar and formulation. After using this tool or service, the author(s) reviewed and edited the content as needed and take(s) full responsibility for the content of the publication.

