## Supplementary figures and images for "Differentiation of Mesenchymal Stem/ Stromal Cells into CD45+ Macrophage-like Cells: Expanding Insights into MSC Plasticity"

### Figure S1

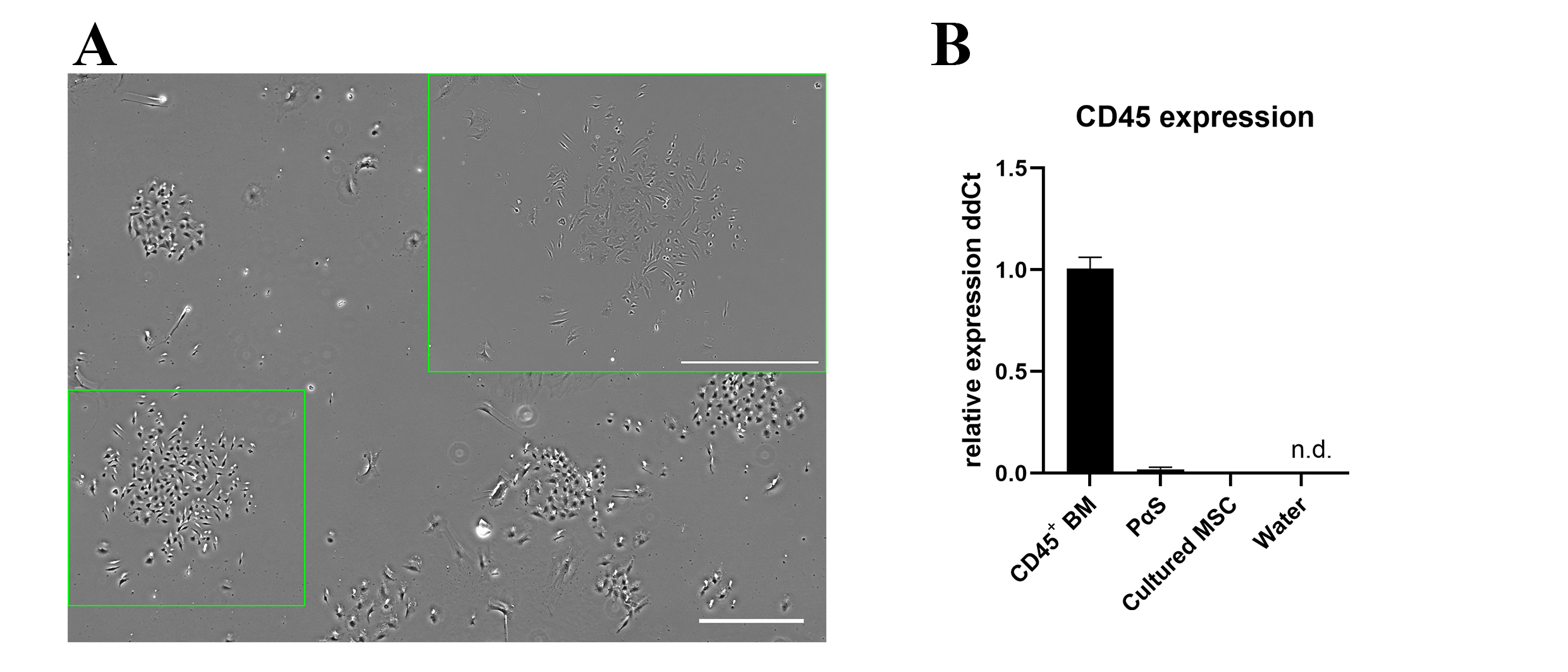

### Figure S2

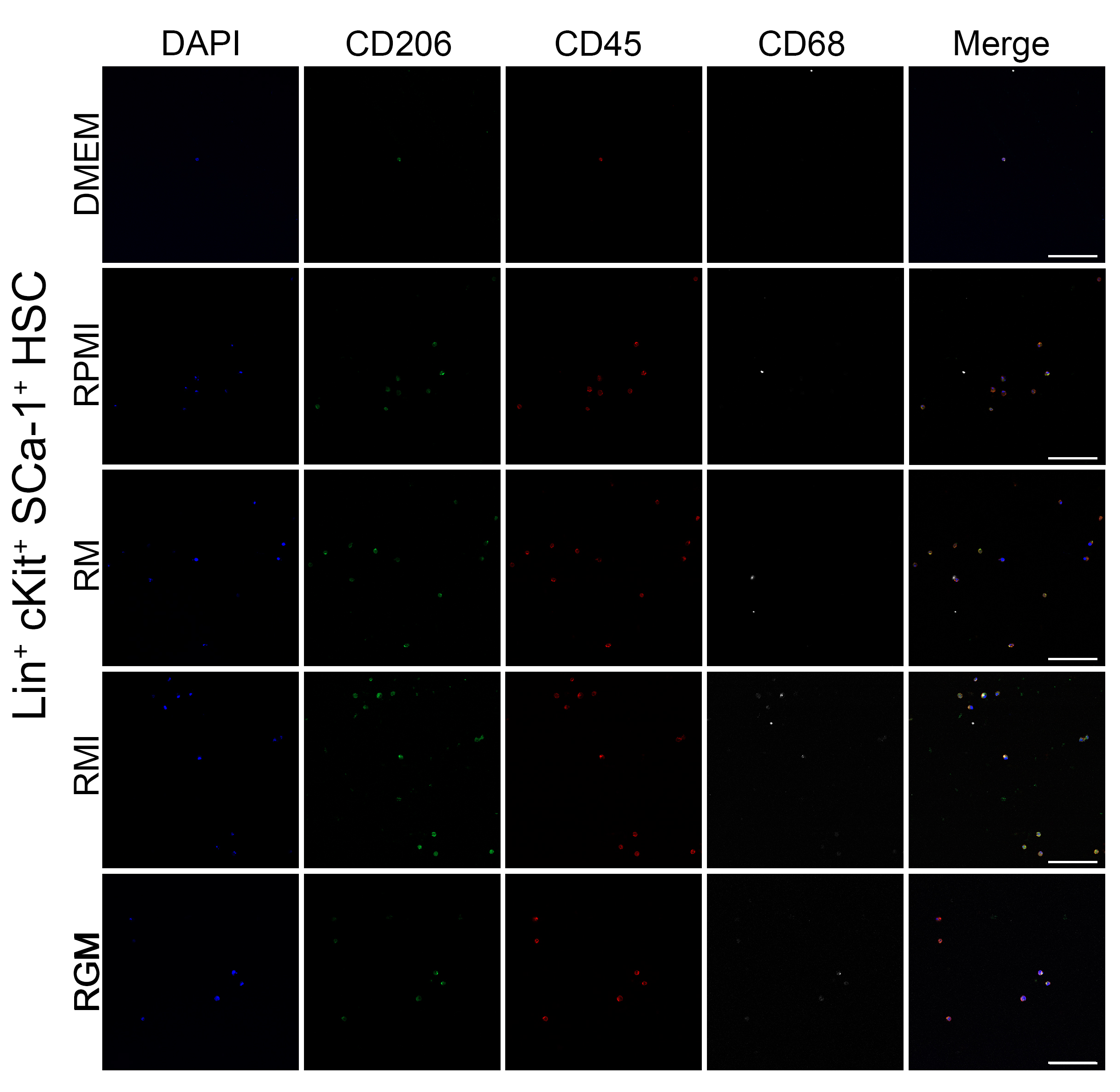
